# Importance of wildlife in the circulation and maintenance of SAT1 and SAT2 foot-and-mouth disease viruses in Africa

**DOI:** 10.1101/2023.05.01.538841

**Authors:** Florian Duchatel, Francois Maree, Louis van Schalkwyk, Barend MdeC Bronsvoort, Samantha Lycett

## Abstract

**Author summary:** Foot and mouth disease (FMD) viruses are endemic in sub-Saharan Africa. Due to the complexity of the disease epidemiology and the lack of available data, there is a need to use phylogenetic approaches to understand the role of potential hosts involved in the circulation and maintenance of the viruses. The uneven host sampling of the available sequences requires us to take advantage of the recent advances in phylogenetic reconstruction. Therefore, using two structural coalescent model approximations we estimated the circulation of FMD virus serotypes SAT1 and SAT2 between cattle, buffalo and impala populations. Our results suggest that in Africa, the impala population seems to act as an intermediate host between the cattle and buffalo populations and play a more important role in the circulation of the viruses than was previously suspected. Until now, the role of the impala population in the circulation of FMDV has been suggested, but never explicitly shown.

## Introduction

Foot and mouth disease (FMD) is a vesicular disease affecting more than 70 species of cloven-hoofed animals, including domestic ruminants and pigs^1^. The causal agent of foot and mouth disease is a positive-sense, single-stranded RNA virus (FMDV). Since the most significant hosts in the natural epidemiology of FMDV are of major importance in the production of food (cattle, sheep, pigs, goats), the disease can potentially lead to important direct and indirect economic impacts in countries with a developed agricultural industry^2–4^.

The most common transmission route between infected and susceptible hosts is by direct contact. In this situation the viral transmission is mechanical, with virus entry through skin cuts or mucosae, following physical contact with infected secretions or excretions^1,5^. Based on the level of cross protection between each strain, we can divide the virus population into seven distinct viral serotypes: O, A, C, Southern African Territories (SAT) 1, SAT 2, SAT 3 and Asia1^6,7^ (N.B. serotype C may be extinct). With five of the seven possible serotypes present in Africa over the last decade and with high regional variances in both their distribution and prevalence, the epidemiology of FMD in Africa is considered to be more complex than anywhere else. It is generally accepted that FMDV originated on the African continent due to the long-term subclinical infection status observed in African Cape buffalo (*Syncerus caffer)* and the important genetic diversity observed in the SAT serotypes^7,8^. However, the rinderpest pandemic largely removed populations susceptible to FMD and, as a result, FMD occurrence declined around the turn of the 20^th^ century, with cases in southern Africa only being reported again in 1931^9^. It is likely that currently circulating lineages of SAT serotypes re-emerged from small numbers of buffalo that survived the rinderpest pandemic once buffalo and livestock numbers had recovered. Moreover, although most FMD outbreaks in sub-Saharan Africa go unrecorded, FMDV is considered endemic in almost all sub-Saharan African countries. This situation might be partially explained by the extensive livestock-raising systems which are practiced in several African regions, leading to an apparent low direct impact from the disease^8^.

Amongst all the wildlife species in Africa that are susceptible to FMDV, only the buffalo and the impala (*Aepyceros melampus*) have been implicated in transmission of the virus to cattle^8,10^. In southern Africa, buffaloes are suspected to be the source for many livestock outbreaks, in particular those concerning the SAT-type FMD viruses for which they are considered the true maintenance host^8,11^. Hence, it is considered that the transmission from buffaloes to domestic animals is the dominant pathway of disease propagation^1,8^. For example, in Ethiopia it has been observed that the cattle populations with the highest FMDV antibody prevalence are those in close contact with wild animals and located near wildlife sanctuaries where large populations of African buffaloes can be found^12,13^ and in Zimbabwe cattle incursions into buffalo areas has been reported as an important driver for FMD outbreaks in livestock^14^. However, there is still a lot of uncertainty regarding the role of buffaloes elsewhere in Africa as a source of livestock outbreaks and how these buffalo populations are able to sustain endemic cycles of infections in livestock^7^. Moreover, although impala do not seem to become persistently infected with FMDV, they have been suspected of acting as intermediate hosts between buffalo and cattle populations in southern Africa^15^.

Due to a lack of a proof-reading mechanism of the RNA-dependent RNA polymerase, replication of the FMDV genome is subject to high mutation rates^6^. The unobserved ecological and population events impacting virus evolution can be estimated while reconstructing the phylogenetic tree of the epidemic^14,16^. By combining genetic and epidemiological information we can estimate the circulation of fast evolving pathogens between multiple discrete states such as the location and the host involved^17,18^.

This approach has already been used on a few occasions to study the role of the cattle and buffalo populations in the circulation and maintenance of FMD viruses in Africa^19,20^. However, the results of such analyses need to be interpreted with caution due to the uneven spatio-temporal sampling across the different discrete states used in these analyses^19,21–23^. It has been shown that the phylogeographical method used in these papers (the “mugration” model^16^) suffers from statistical bias, exacerbated by an uneven sampling between the populations^24,25^. A way to mitigate the impact of these biases is to use a structural coalescent phylogenetic model approximation^24,26^. Such approaches, by taking into account the effect of the migration events on the phylogeny structure, have a better model specification in their phylogenetic and evolutionary parameter estimations such as the effective population size (Ne) of the viral populations estimated in each of the analyzed discrete states^27^. Ne being the size of an idealized population randomly mating and having the same gene frequency changes as the whole population under study^28^.

The aim of this paper is to compare the results of two recent structural coalescent model approximations (BASTA^27^ and MASCOT^29^) with the previously used “mugration” approach. Therefore, we estimated the evolution and transmission between cattle, impala and buffalo populations of FMDV serotypes SAT1 and SAT2 across Africa.

## Materials and methods

We used two datasets of FMDV SAT1 and SAT2 sequences from a previous analysis^30^. The FMDV SAT1 dataset was composed of 117 sequences with dates ranging from 1961 to 2015, including 16 sequences from impalas, 35 sequences from buffaloes and 68 sequences from cattle. The FMDV SAT2 dataset was composed of 135 sequences with dates ranging from 1970 to 2015, including 7 sequences from impalas, 34 sequences from buffaloes and 98 sequences from cattle (see Table 1 and Table 2) for the number of sequences per host and location and Supplementary Figure S1 and Supplementary Figure S2 for the annotated ‘mugration’ trees with the host and location for each sequence).

**Table 1.**
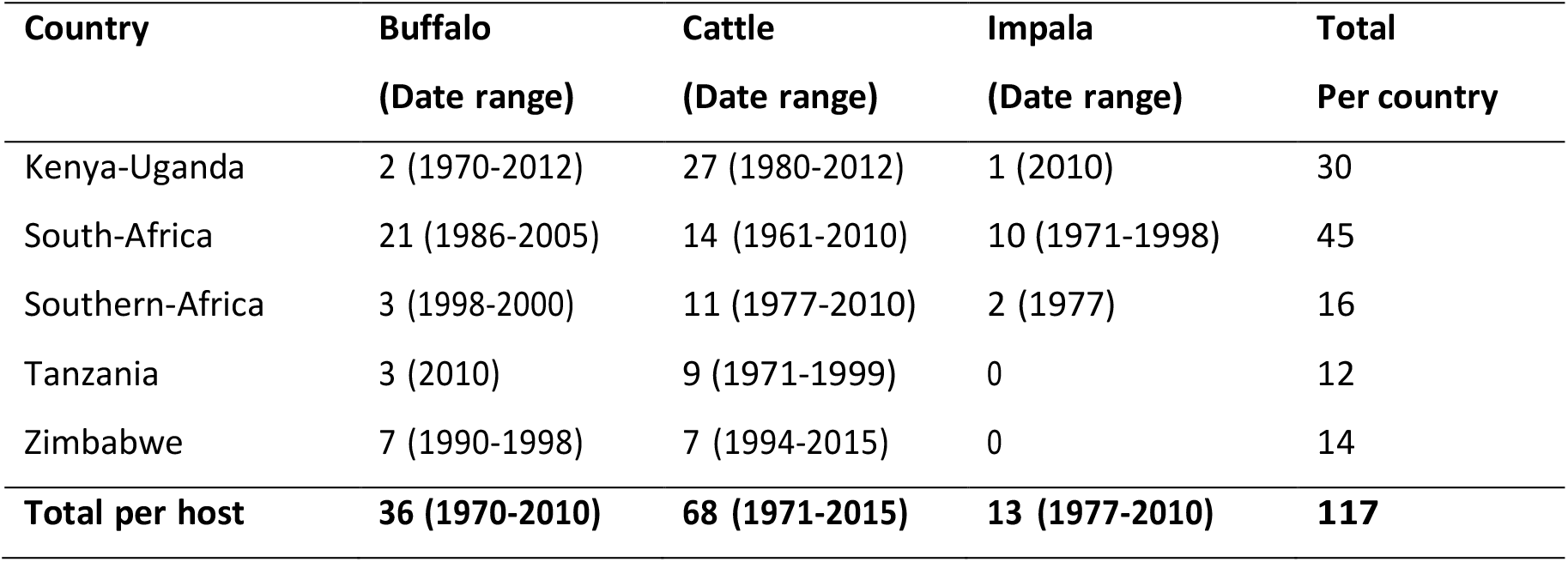
Host and origin of the SAT1 sequences utilized in the phylogenetical analysis.The year of sampling range can be seen for each one of the combinations of host and origin.

**Table 2.**
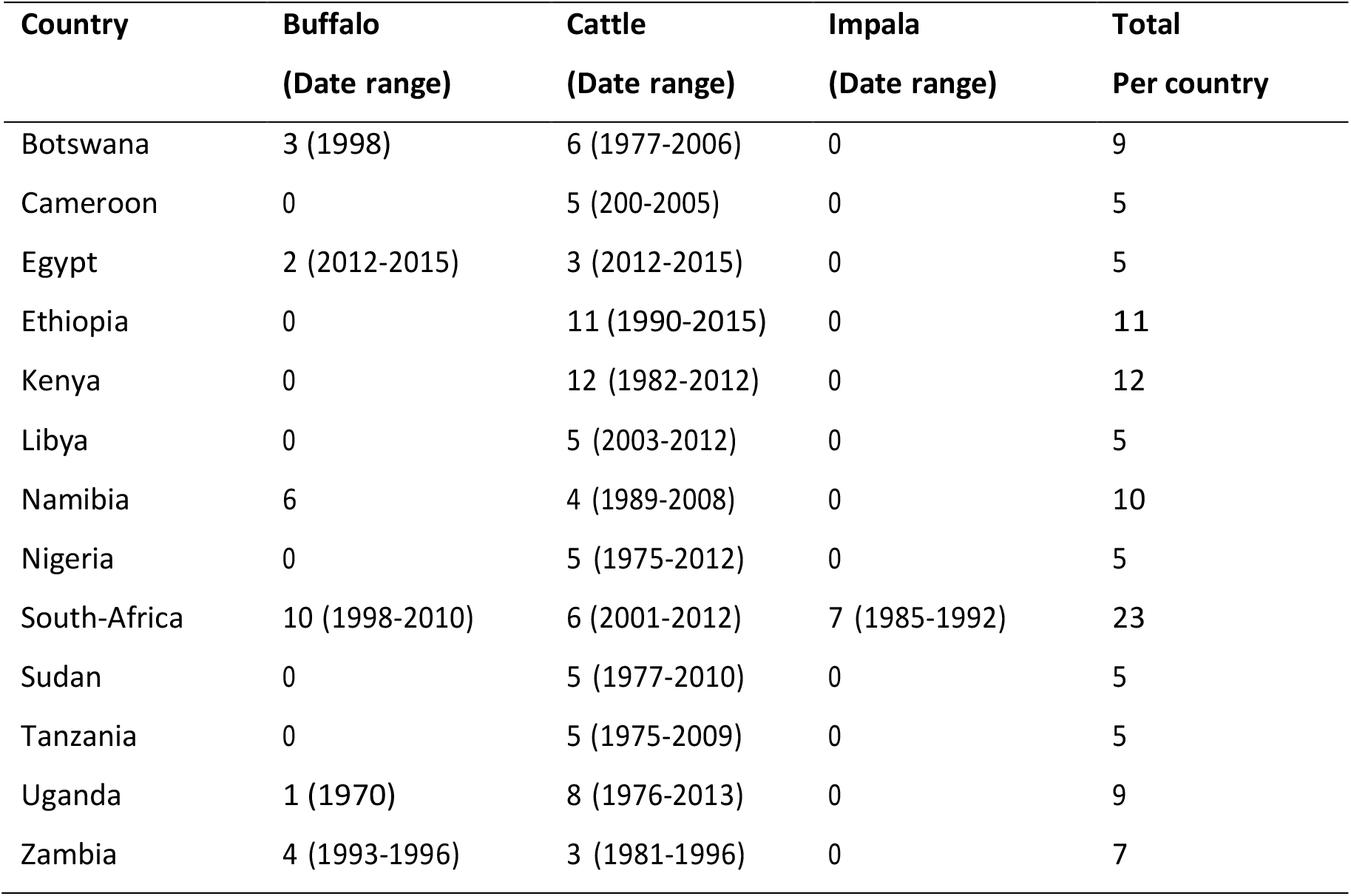
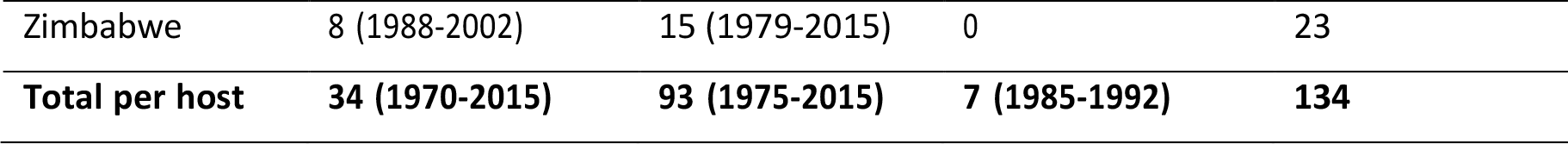
Host and origin of the SAT2 sequences utilised in the phylogenetical analysis. The year of sampling range can be seen for each one of the combinations of host and origin.

The “mugration” phylogenetic trees were reconstructed using BEAST 1.8 with the BEAGLE library^31^. For both serotypes, a Hasegawa-Kishono–Yano (HKY) nucleotide substitution model with a constant clock model and a Bayesian skygrid population model were chosen to model the evolution the virus^32,33^.

We first reconstructed the time-scaled phylogenetic trees for the two studied serotypes by combining at least two independent Markov Chain Monte Carlo runs of 40 million steps sampling every thousand with a 10% burn-in. Thereafter, to reduce the computation time needed for the hosts diffusion analysis, we used subsets of 1000 trees from the original posterior distributions of trees as empirical tree distributions.

Simultaneously, we performed a phylogenetic tree reconstitution using two structural coalescent model 82approximations with the BASTA^26^ and MASCOT^28^ packages available within BEAST^34^. For both models and serotypes, we used an HKY model of molecular evolution and a constant clock model. For each analysis we combined three converging runs of at least 50 million steps (BASTA) or 10 million steps (MASCOT). For all the analysis we used TreeAnnotator to summarize maximum clade credibility (MCC) trees and FigTree version 1.4.1 to visualize the annotated trees^34,35^.

For the ‘mugration’ and BASTA approaches we were able to perform a Markov jump analysis to determine the number of transmission events that occurred between the three host over the whole phylogeny^20^.

## Results

The SAT1 serotype clock rate estimates were similar at around 2×10^−3^ substitution/site/year for the three phylogenetic methods. We observed a tree height of 260 years for the “mugration” method, and a tree height closer to 200 years for the two structural coalescent approximation methods. Both BASTA and MASCOT models estimated that an important viral population present in buffalo (736 ± 117 for MASCOT and 903 ± 187.6 for BASTA) and a medium viral population size present in cattle (7.35 ± 4.6 for BASTA and 4.19 ± 3.14 for MASCOT) had a role in the serotype transmission. The two models estimated that only a small viral population present in impala was involved in the circulation of the disease (2.19 ± 1.15 for BASTA and 2.43 ± 1.45 for MASCOT) (see Supplementary table S1 and Supplementary figure S3).

In all three methods we can observe the same three main clades. With two out of the three clades of the reconstructed tree being almost entirely composed of cattle nodes (clades 1 and 3), the “mugration” approach estimated an important role for cattle in the transmission of the SAT1 virus (see Figure 1a). With the “mugration” model we observe that in only a few occasions (mostly in clade 2) the impala population acted as an intermediate host between the buffalo and cattle populations. However, for both structural coalescent models, we observed that most of the reconstructed trees were either composed of buffalo nodes (for BASTA) or a mixture of buffalo and impala nodes (for MASCOT), with the cattle population being present only at the tips of the trees (see Figure 1b and Figure 1c). In both structural approaches we observe multiple occasions where the impala population acted as the intermediate host between the buffalo and cattle populations. The Markov jump analysis results and the estimated transmission rates between the different populations for FMDV serotype SAT1 emphases the observed differences between the “mugration” model and the two structural coalescent approximation approaches.

**Figure 1.**
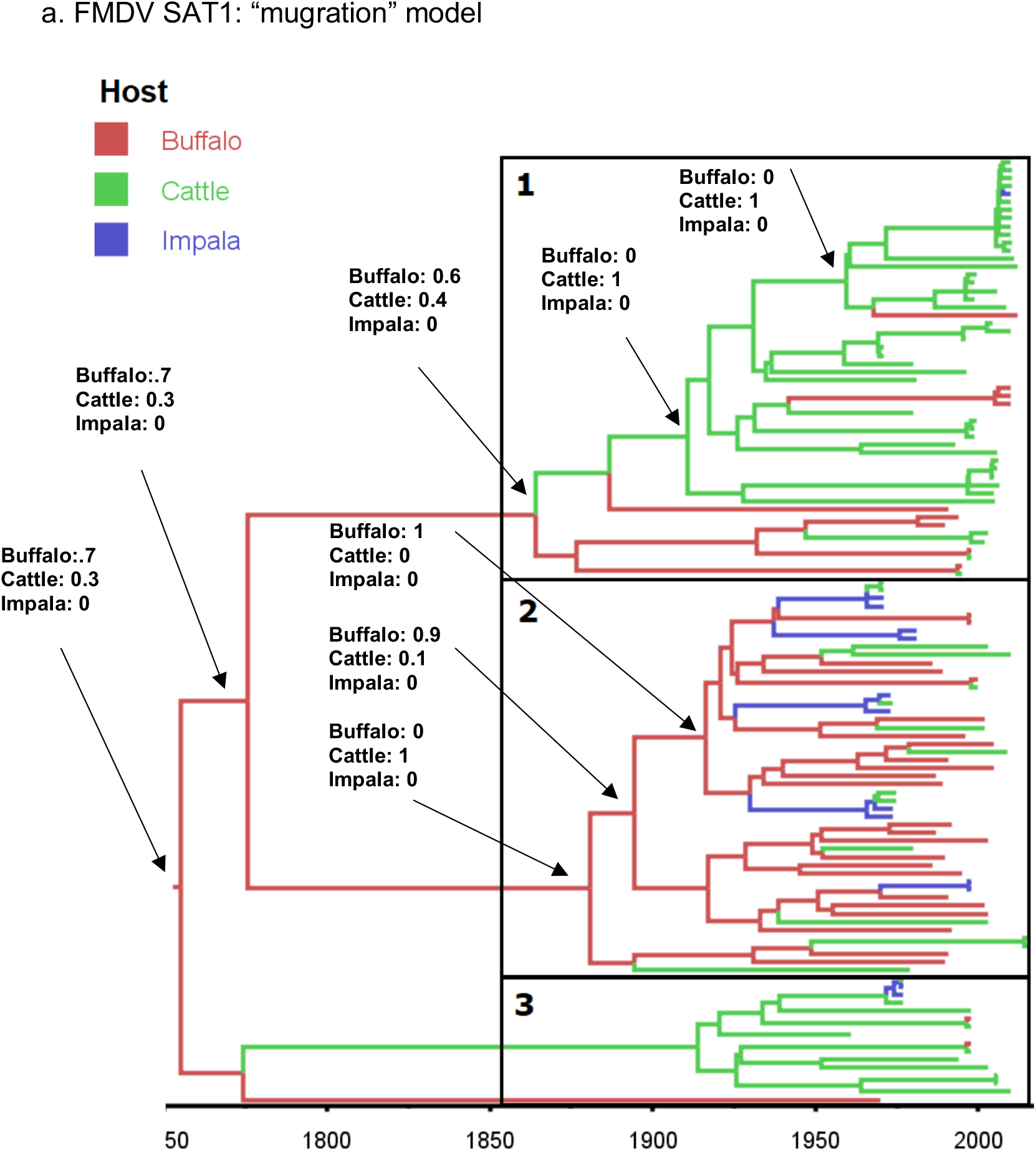
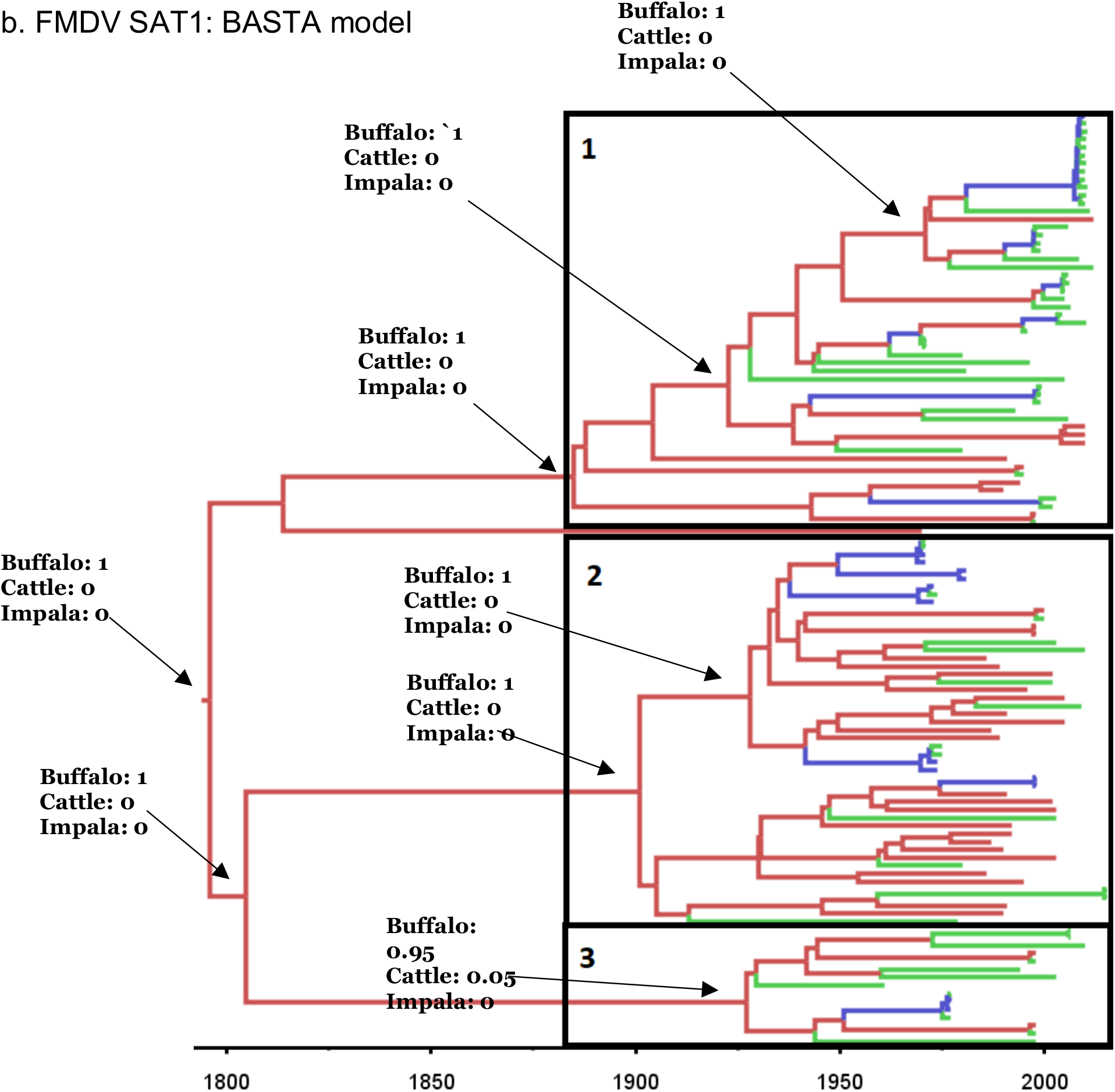
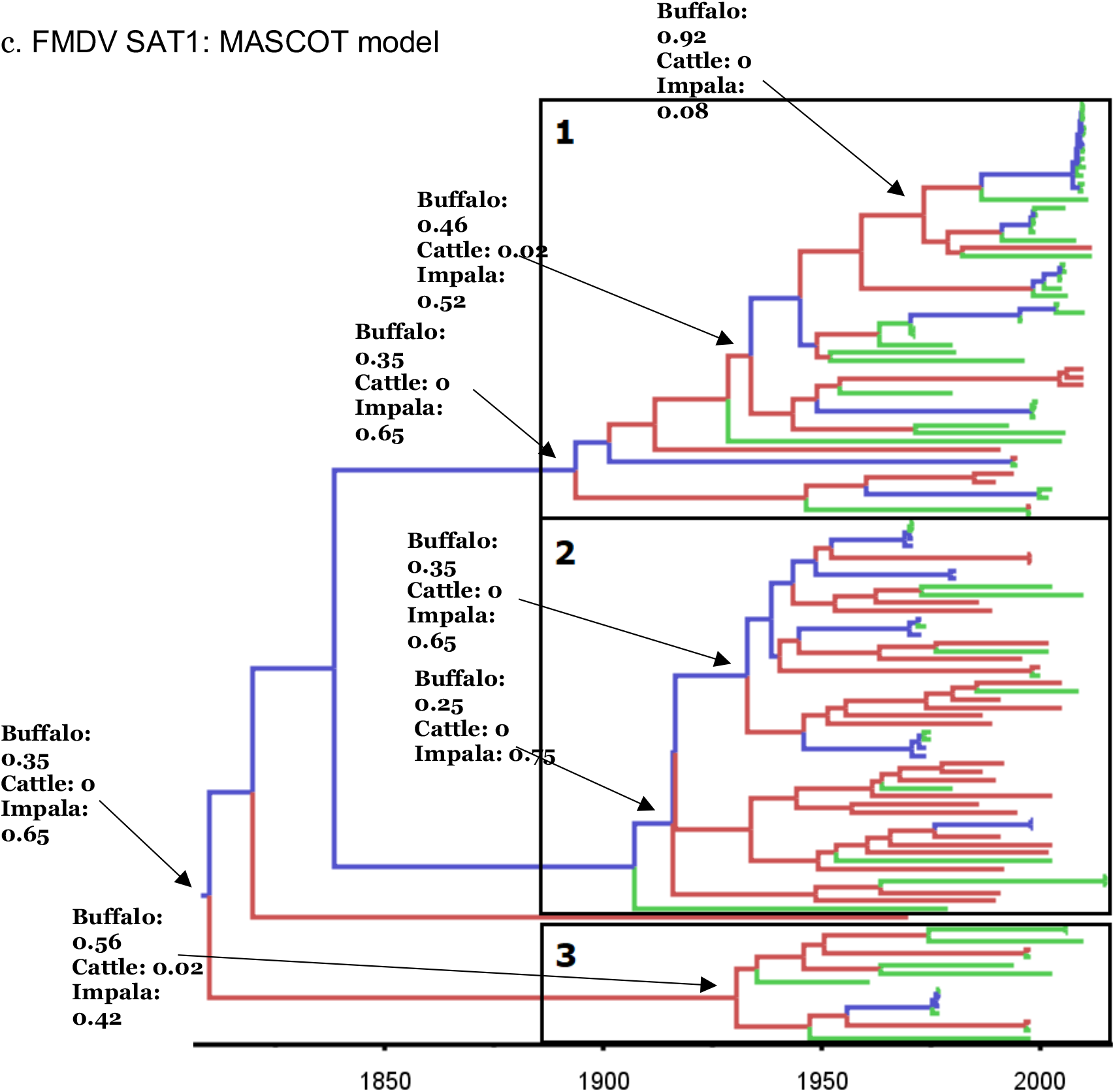
Bayesian MCC time scaled discrete phylogeographic tree for the serotype SAT1 using 113 VP1 sequences. a. Phylogenetic tree estimated using the “mugration” approach implemented in BEAST b. Phylogenetic tree estimated using the BASTA approach implemented in BEAST2. c. Phylogenetic tree estimated using the MASCOT approach implemented in BEAST2. The phylogeny branches are coloured according to their descendent nodes host with the key for colours shown on the upper left of the figure. The identified clades were isolated and numerated. Specific nodes of the trees were annotated with hosts posterior probabilities.

For both structural coalescent models, we obtained similar transmission rates between the three populations (see Figure 2b and Figure 2c). We estimated low transmission rates from the impala and cattle populations toward the buffalo population (less than 1-2). We observed high transmission rates from the impala and buffalo population toward the cattle population (1.59 ± 0.64 and 1.56 ± 1.42 for BASTA and 1.27 ± 0.39 and 0.93 ± 0.44 for MASCOT) and lower transmission rates from the buffalo and cattle populations to the impala population (0.6 ± 0.32 and 0.46 ± 0.43 for BASTA and 0.26 ± 0.18 and 0.33 ± 0.39 for MASTCOT). These results are quite different to those obtained with the “mugration” model (see Figure 2a). The main differences with the structural approaches being the high transmission rates estimated between the buffalo and cattle populations (respectively 1.9 ± 0.97 and 1.18 ± 0.69) and the transmission rate of 0.35 ± 0.35 from the impala to the buffalo population.

**Figure 2.**
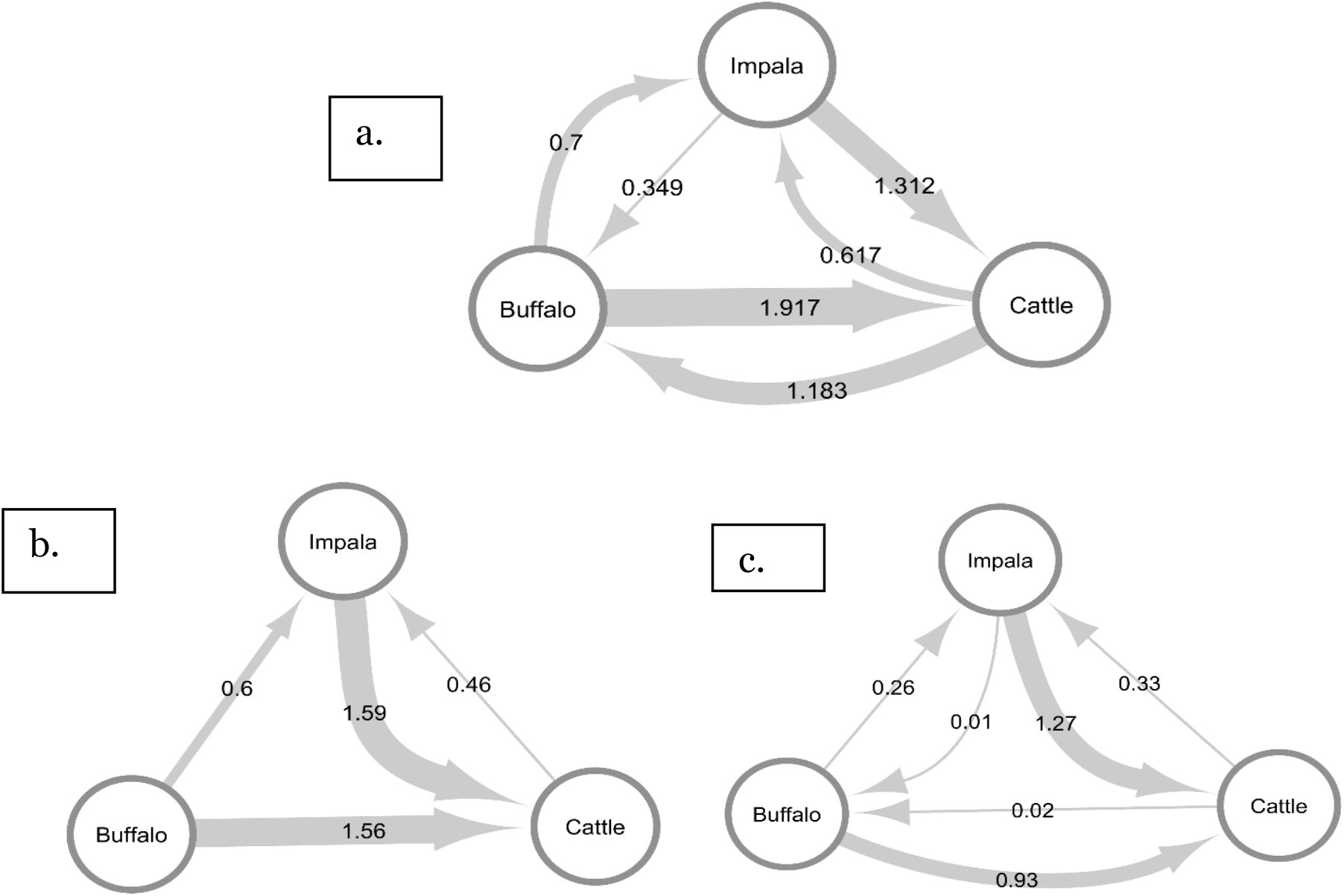
Estimated transmission rates between the three potential hosts for the SAT1 FMDV serotype. a. Using the “Mugration” approach in BEAST. b. Using the BASTAapproach in BEAST2. c. Using the MASCOT approach in BEAST2.

Using the “mugration” model, we determined numerous transmission events between buffalo and cattle populations and lower numbers of transmission events between the impala and cattle populations (see Figure 3a). In comparison, using BASTA, we observed few transmission events from the cattle to the buffalo populations but more transmission events between the impala and cattle populations (see Figure 3b).

**Figure 3.**
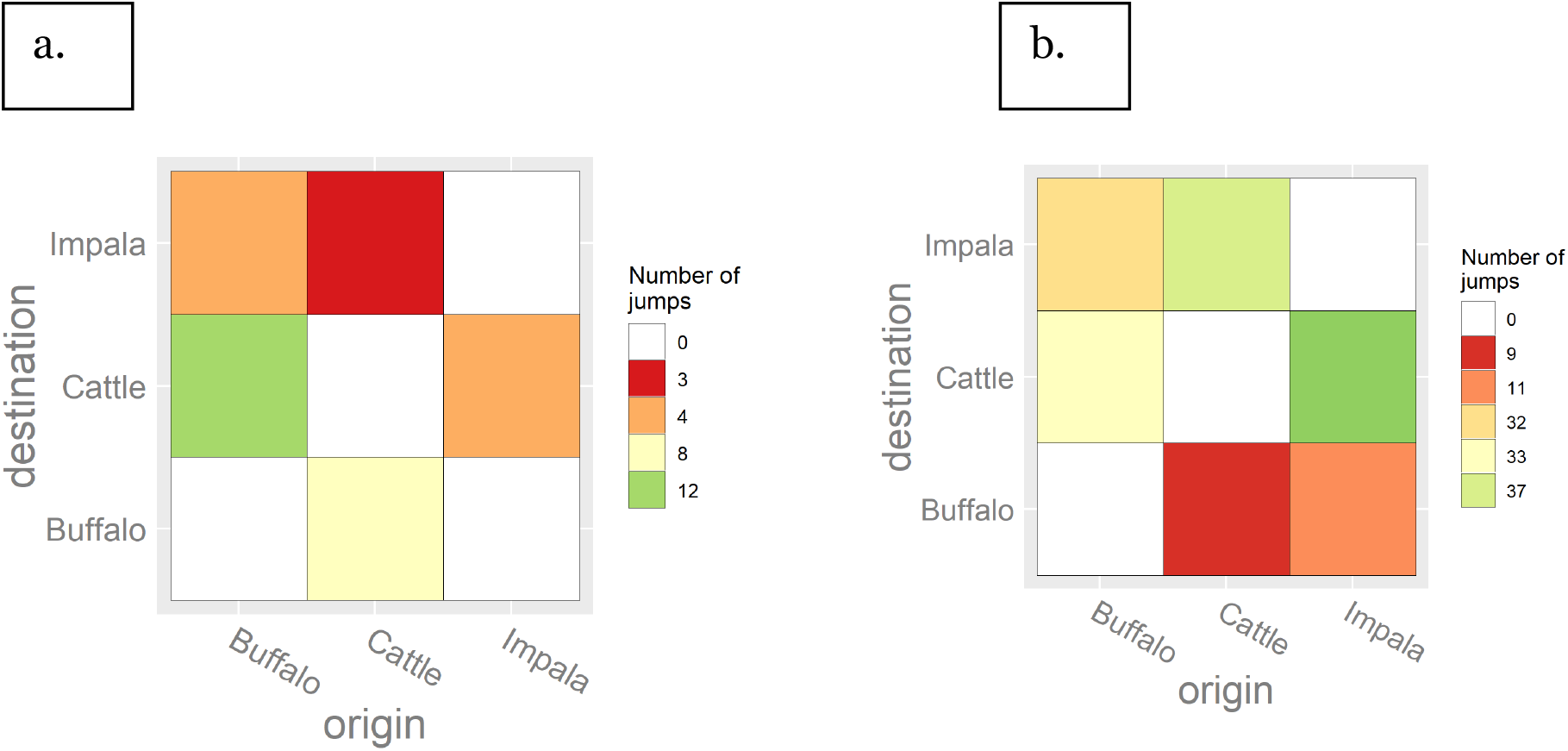
Heatmap showing the number of transitions between the sampled hosts for the SAT1 FMDV serotype obtained through a Markov jump analysis. The heatmaps are coloured according to the number of estimated transitions between hosts. a. Host transition using the”mugration” model b. Host transition using the BASTA model.

When analyzing the FMDV SAT2 serotype dataset we estimated a similar clock rate of 1.1×10^−3^ substitution/site/year between the tree models. However, we estimated a longer tree height of 430 years for the “mugration” method than for the two structural coalescent approximation methods that had a tree height closer to 350 years. Both BASTA and MASCOT models estimated a similar buffalo population size (around 1200 for both models) and impala population size (2/3 in both models) involved in the SAT2 serotype spread. However, the BASTA model estimated a slightly larger cattle population (3.9 ± 1.89) than MASCOT (1.8 ± 1) (see Supplementary table **S1** and Supplementary table **S2)**

With all three methods we isolated the same fives main clades. Because the “mugration” model estimated that two out of five clades composing the phylogenetic tree of SAT2 were entirely composed of cattle nodes (clades two and five), it estimated an important role of the cattle population in the transmission of the virus (see Figure 4a). However, for this serotype we cannot observe transmission events from the impala toward the cattle populations. In comparison, with most of the tree composed of buffalo nodes, the BASTA and MASCOT models estimate a less important role for the cattle population in the transmission of FMDV SAT2 serotype (see Figure 4b and Figure 4c.).

**Figure 4.**
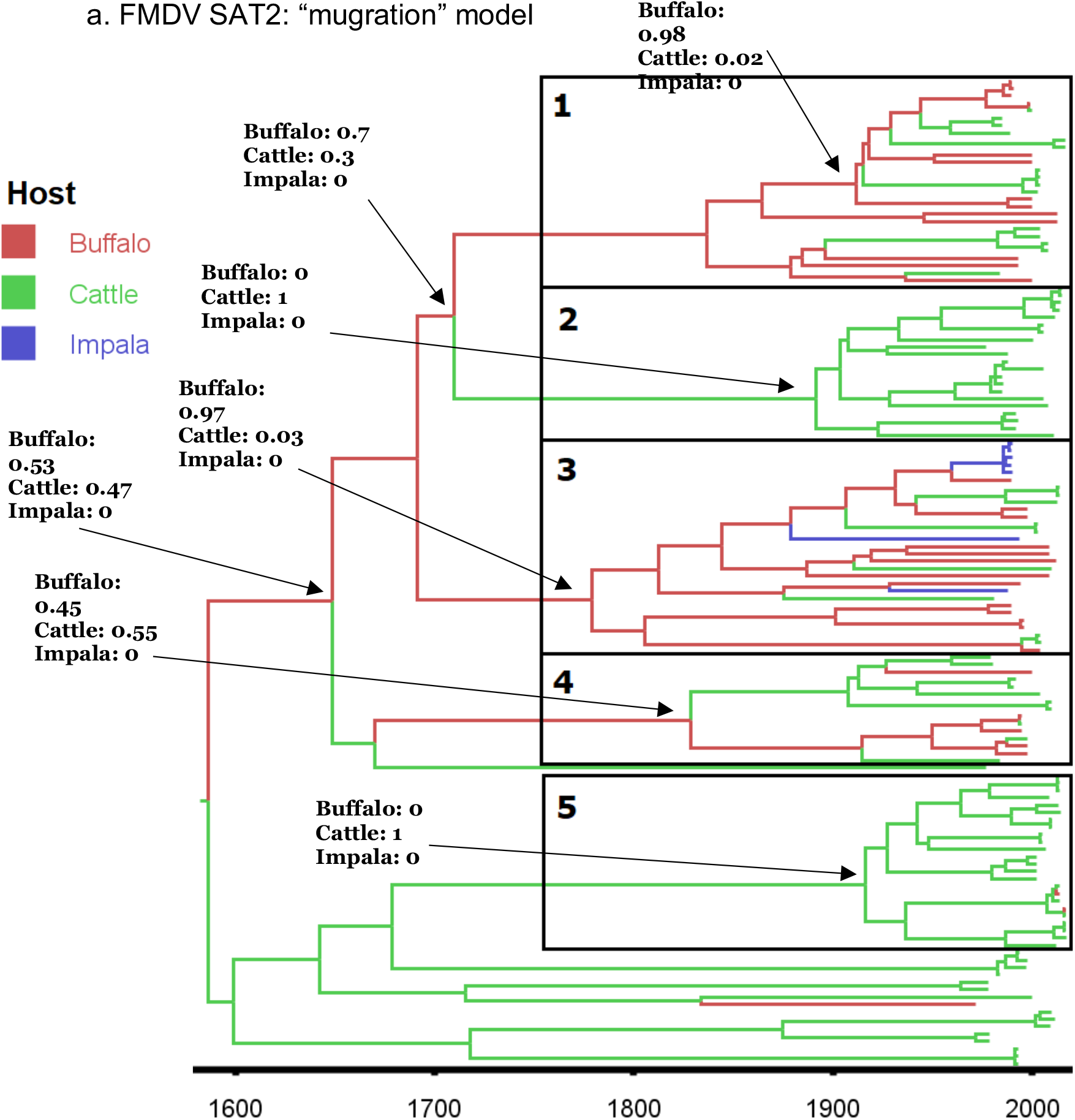
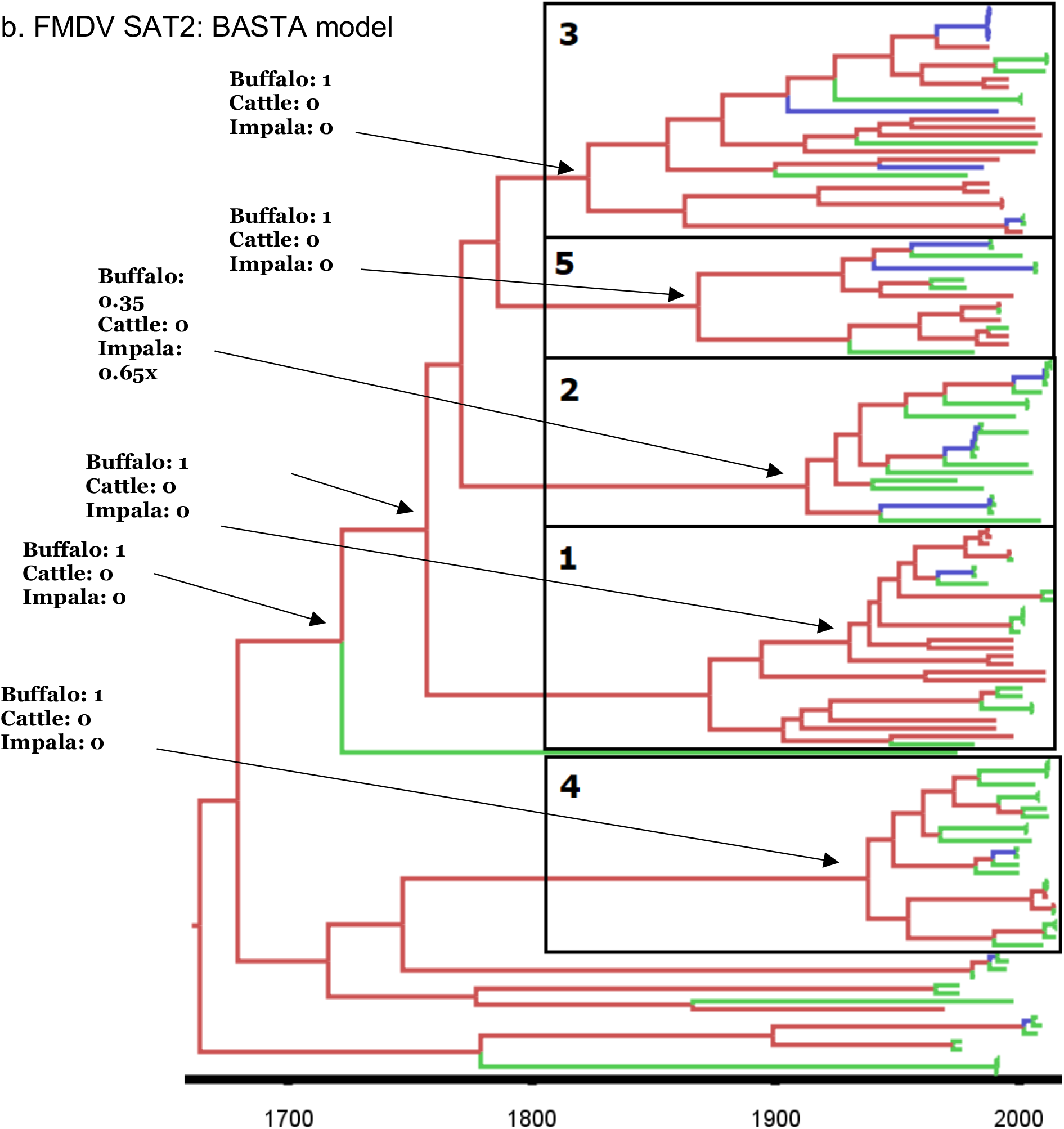
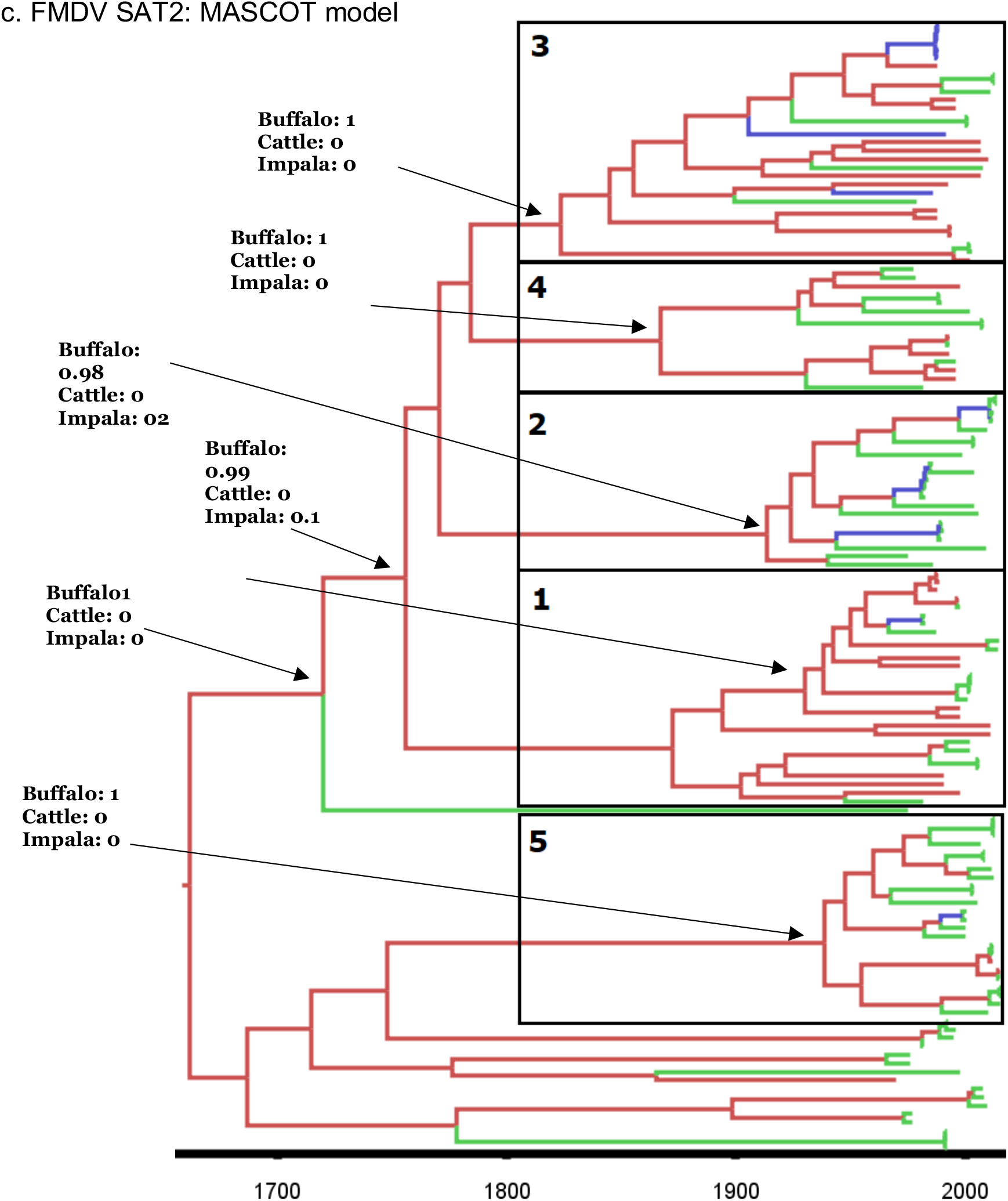
Bayesian MCC time scaled discrete phylogeographic tree for the serotype SAT2using 135 VP1 sequences. a. Phylogenetic tree estimated using the “mugration” approach implemented in BEAST b. Phylogenetic tree estimated using the BASTA approach implemented in BEAST2. c. Phylogenetic tree estimated using the MASCOTapproach implemented in BEAST2. The phylogeny branches are coloured according totheir descendent nodes host with the key for colours shown on the upper left of the figure. The identified clades were isolated and numerated. Specific nodes of the trees were annotated with hosts posterior probabilities

Additionally, in these two models we estimated multiple transmission events of FMDV SAT2 serotypes from buffalo to impala and then to the cattle population. Again, as with the FMDV SAT1 serotype, the estimated number of transitions and transmission rates between the different populations estimated for the SAT2 serotype stresses the differences between the “mugration” model and both structural coalescent approximation approaches.

In both the BASTA and MASCOT models, we estimated low transmission rates (less than 1-2) from the impala and cattle populations toward the buffalo population (see Figure 5b and Figure 5c). In both approaches, we estimated higher rates of transmission from buffaloes to impalas (1 ± 0.44 in BASTA and in 0.466 ± 0.28 in MASCOT) than from cattle to impalas (0.24 ± 0.31 in BASTA and 0.25 ± 0.33 in MASCOT). By looking at the transmission rate, the main difference between the two structural coalescent models is that the BASTA model estimated a higher transmission rate from impalas to cattle than from buffaloes to cattle (1.38 ± 0.77 and 0.7 ± 0.46), while the MASCOT model estimated similar transmission rates (0.95 ± 0.66 and 0.96 ± 0.43). In comparison to the structural coalescent approximation approaches, most of the transmission rates obtained with the “mugration” model were between 0.5 and 1, except for the transmission rate from the buffalo to the cattle population which is equal to 2.8 ± 1.44 (see Figure 5a).

**Figure 5.**
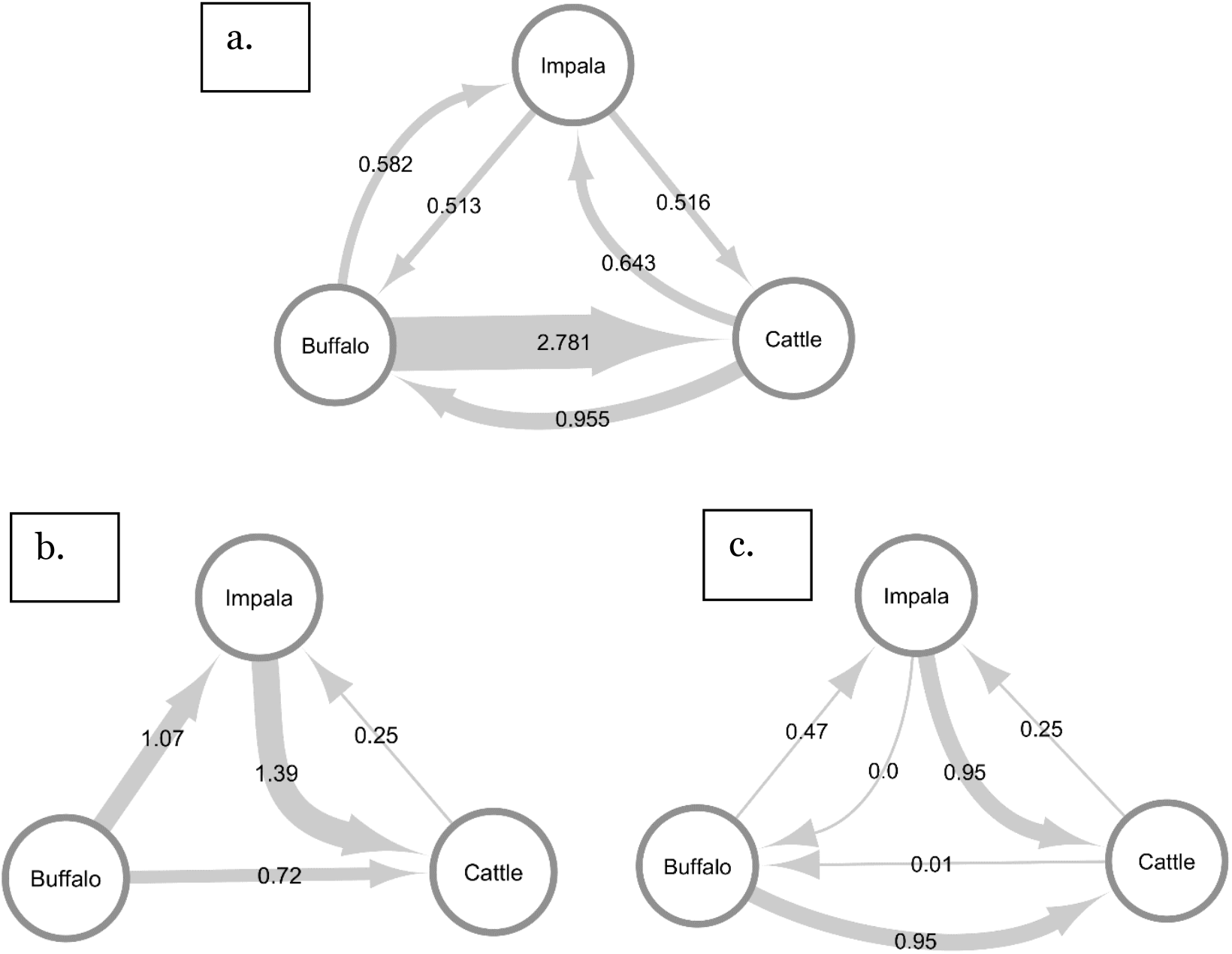
Estimated transmission rates between the three potential hosts for the SAT2 FMDV serotype. a. Using the “Mugration” approach in BEAST. b. Using the BAST Aapproach in BEAST2. c. Using the MASCOT approach in BAST2.

The “mugration” model estimated that multiple transmission events occurred between the buffalo and cattle populations and that no transmission events occurred between the impala and cattle populations (see Figure 6a). In comparison, the BASTA approach estimated multiple transmission events from the impala and buffalo populations toward the cattle population and a small number of transmissions with the cattle population as origin (see Figure 6b).

**Figure 6.**
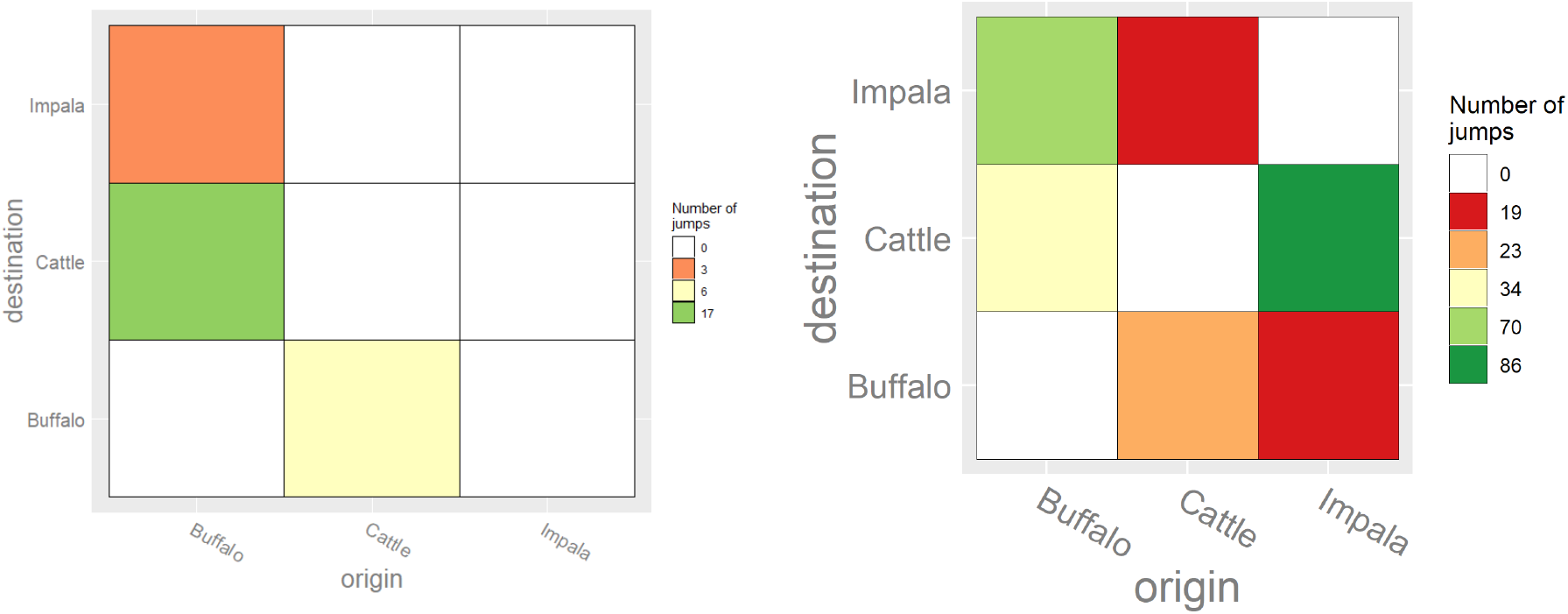
Heatmap showing the number of transitions between the sampled hosts for theSAT2 FMDV serotype. The heatmaps are colored according to the number of estimated transitions between hosts. a. Host transition using the “mugration” model b. Host transition using the BASTA model.

## Discussion

In this study, we applied three recently developed methods of phylogenetic reconstruction to estimate the transmission and circulation of the VP1 gene sequence between cattle, buffalo and impala populations for two FMDV serotypes (SAT1 and SAT2) in Africa. Two of these methods, the BASTA and MASCOT models, consider the existence of different host populations to reconstruct the evolution of the pathogen. The last approach, the “mugration” model, does not consider the existence of multiple host populations, making it more prone to statistical errors caused by sampling bias. Considering that both our BASTA and MASCOT models produced similar estimates, it seems that the host structure of the populations plays an important role in the outcome of the analyses. However, the differences observed between the “mugration” and the BASTA/MASCOT models were expected, given the incomplete nature of the sampling for the two serotypes^26^.

Using the “mugration” model, we estimated an important role for the buffalo and cattle populations in the maintenance of both FMDV SAT1 and SAT2 viruses in Africa. The model produced high transmission rate estimates between the two populations and much lower rates for transmission to impala suggesting the impala population acts more as a spill over host for the viruses. With this strong connection between cattle and buffalo populations and the apparent lesser role played by the impala population in the circulation of the viruses, the “mugration” model results are consistent with those made in previous studies^21,22^.

However, the BASTA and MASCOT models, suggest a different interaction between the three hosts in the circulation and maintenance of the viruses. The higher number, proportionally, of transmission events, as well as higher transmission rates, between buffalo and impala populations produced by the BASTA model compared to the “mugration” approach is consistent with the real-life observations of regular transmission events between buffaloes and impalas^13,16^. Moreover, the observed lower transmission rates toward the buffalo population and their large population sizes in both structural approaches, reinforce the general consensus that buffaloes are the maintenance and original host of the SAT serotypes^8^.

Additionally, although the estimated impala population size was small, the large transmission rate from the impala population toward the cattle population, estimated by the structural approaches, suggest an at least as important role played by the impala population in the viral transmission toward the cattle population in southern Africa. Overall, in both structural approaches, we observed that the impala population plays the role of an intermediate host between the buffalo and cattle populations, with the cattle as final host and the buffalo as original host of the viruses.

Overall, our results point out a more important role of the impala population than what was previously thought with buffaloes as being the original host of the virus. However, since most of the analyzed samples originate from southern Africa, our observations could be the result of locally observed relationships between the three populations and might not be generalized to sub-Saharan Africa. This situation emphasizes the need of a more systematic sampling of impala and buffalo populations. Such approach would enable us to detect currently unrecorded/sub-clinical buffalo/impala outbreaks and confirm the results of this analysis more globally. Similar to what the “mugration” approach suggests, our current knowledge sees impalas as intermediate hosts whereas the structural coalescent results suggest a simultaneous buffalo to cattle and buffalo to impala transmission, with a possible final impala to cattle transmission. Our study is the first phylogenetic analysis using a structural coalescent approach to reconstruct the circulation and maintenance of the SAT serotypes between multiple hosts. It is also the first time that the importance of the impala population in the transmission and circulation of the virus has been quantified.

The fact that the SAT serotypes are mainly maintained within wild buffalo populations does not go against the observation that most of the recent spread of the viruses in Africa seems to be driven by livestock movement^36–38^. Moreover, this observation is in concordance with the theory that currently circulating lineage of SAT serotypes re-emerged from a small number of wild buffaloes that survived the African rinderpest epidemic of 1975^10^. Although the use of more complex phylogenetic models allows us to gain more insight into the epidemiology and evolution of FMDVs, greater sampling effort is needed to obtain more diversity in the types of hosts sampled and the temporal and spatial distributions of isolates in order to observe and explain these results epidemiologically. The use of analytical methods is important in uncovering the drivers of disease maintenance and spread, and thus will help in developing a modern approach to FMD control where different hosts could be targeted and controlled in different regions.

## Acknowledgments

SL and MB are supported by an Institute Strategic Programme Grant (XXXX), ???

