## Supporting Material for "Importance of wildlife in the circulation and maintenance of SAT1 and SAT2 foot-and-mouth disease viruses in Africa"

**Table 1.** Host and origin of the SAT1 sequences utilized in the phylogenetical analysis.The year of sampling range can be seen for each one of the combinations of host and origin.

| **Country** | **Buffalo (Date range)** | **Cattle (Date range)** | **Impala (Date range)** | **Total Per country** |
|---|---|---|---|---|
| Kenya-Uganda | 2 (1970-2012) | 27 (1980-2012) | 1 (2010) | 30 |
| South-Africa | 21 (1986-2005) | 14 (1961-2010) | 10 (1971-1998) | 45 |
| Southern-Africa | 3 (1998-2000) | 11 (1977-2010) | 2 (1977) | 16 |
| Tanzania | 3 (2010) | 9 (1971-1999) | 0 | 12 |
| Zimbabwe | 7 (1990-1998) | 7 (1994-2015) | 0 | 14 |
| **Total per host** | **36 (1970-2010)** | **68 (1971-2015)** | **13 (1977-2010)** | **117** |

**Table 2.** Host and origin of the SAT2 sequences utilised in the phylogenetical analysis.The year of sampling range can be seen for each one of the combinations of host and origin.

| **Country** | **Buffalo (Date range)** | **Cattle (Date range)** | **Impala (Date range)** | **Total Per country** |
|---|---|---|---|---|
| Botswana | 3 (1998) | 6 (1977-2006) | 0 | 9 |
| Cameroon | 0 | 5 (200-2005) | 0 | 5 |
| Egypt | 2 (2012-2015) | 3 (2012-2015) | 0 | 5 |
| Ethiopia | 0 | 11 (1990-2015) | 0 | 11 |
| Kenya | 0 | 12 (1982-2012) | 0 | 12 |
| Libya | 0 | 5 (2003-2012) | 0 | 5 |
| Namibia | 6 | 4 (1989-2008) | 0 | 10 |
| Nigeria | 0 | 5 (1975-2012) | 0 | 5 |
| South-Africa | 10 (1998-2010) | 6 (2001-2012) | 7 (1985-1992) | 23 |
| Sudan | 0 | 5 (1977-2010) | 0 | 5 |
| Tanzania | 0 | 5 (1975-2009) | 0 | 5 |
| Uganda | 1 (1970) | 8 (1976-2013) | 0 | 9 |
| Zambia | 4 (1993-1996) | 3 (1981-1996) | 0 | 7 |

| | | | | |
|---|---|---|---|---|
| Zimbabwe | 8 (1988-2002) | 15 (1979-2015) | 0 | 23 |
| **Total per host** | **34 (1970-2015)** | **93 (1975-2015)** | **7 (1985-1992)** | **134** |

The “mugration” phylogenetic trees were reconstructed using BEAST 1.8 with the BEAGLE library[31]. For both serotypes, a Hasegawa-Kishono–Yano (HKY) nucleotide substitution model with a constant clock model and a Bayesian skygrid population model were chosen to model the evolution the virus[32,33].

We first reconstructed the time-scaled phylogenetic trees for the two studied serotypes by combining at least two independent Markov Chain Monte Carlo runs of 40 million steps sampling every thousand with a 10% burn-in. Thereafter, to reduce the computation time needed for the hosts diffusion analysis, we used subsets of 1000 trees from the original posterior distributions of trees as empirical tree distributions.

Simultaneously, we performed a phylogenetic tree reconstitution using two structural coalescent model approximations with the BASTA[26] and MASCOT[28] packages available within BEAST[34]. For both models and serotypes, we used an HKY model of molecular evolution and a constant clock model. For each analysis we combined three converging runs of at least 50 million steps (BASTA) or 10 million steps (MASCOT). For all the analysis we used TreeAnnotator to summarize maximum clade credibility (MCC) trees and FigTree version 1.4.1 to visualize the annotated trees[34,35].

For the ‘mugration’ and BASTA approaches we were able to perform a Markov jump analysis to determine the number of transmission events that occurred between the three host over the whole phylogeny[20].

### Results

The SAT1 serotype clock rate estimates were similar at around $2x10^{-3}$ substitution/site/year for the three phylogenetic methods. We observed a tree height of 260 years for the “mugration” method, and a tree height closer to 200 years for the two structural coalescent approximation methods. Both BASTA and MASCOT models estimated that an important viral population present in buffalo ($736 \pm 117$ for MASCOT and $903 \pm 187.6$ for BASTA) and a medium viral population size present in cattle ($7.35 \pm 4.6$ for BASTA and $4.19 \pm 3.14$ for MASCOT) had a role in the serotype transmission. The two models estimated that only a small viral population present in impala was involved in the circulation of the disease (2.19 $\pm$ 1.15 for BASTA and 2.43 $\pm$ 1.45 for MASCOT) (see Supplementary table S1 and Supplementary figure S3).

In all three methods we can observe the same three main clades. With two out of the three clades of the reconstructed tree being almost entirely composed of cattle nodes (clades 1 and 3), the "mugration" approach estimated an important role for cattle in the transmission of the SAT1 virus (see Figure 1a). With the "mugration" model we observe that in only a few occasions (mostly in clade 2) the impala population acted as an intermediate host between the buffalo and cattle populations. However, for both structural coalescent models, we observed that most of the reconstructed trees were either composed of buffalo nodes (for BASTA) or a mixture of buffalo and impala nodes (for MASCOT), with the cattle population being present only at the tips of the trees (see Figure 1b and Figure 1c). In both structural approaches we observe multiple occasions where the impala population acted as the intermediate host between the buffalo and cattle populations. The Markov jump analysis results and the estimated transmission rates between the different populations for FMDV serotype SAT1 emphases the observed differences between the "mugration" model and the two structural coalescent approximation approaches.

For both structural coalescent models, we obtained similar transmission rates between the three populations (see Figure 2b and Figure 2c). We estimated low transmission rates from the impala and cattle populations toward the buffalo population (less than 1-2). We observed high transmission rates from the impala and buffalo population toward the cattle population (1.59 $\pm$ 0.64 and 1.56 $\pm$ 1.42 for BASTA and 1.27 $\pm$ 0.39 and 0.93 $\pm$ 0.44 for MASCOT) and lower transmission rates from the buffalo and cattle populations to the impala population (0.6 $\pm$ 0.32 and 0.46 $\pm$ 0.43 for BASTA and 0.26 $\pm$ 0.18 and 0.33 $\pm$ 0.39 for MASTCOT). These results are quite different to those obtained with the "mugration" model (see Figure 2a). The main differences with the structural approaches being the high transmission rates estimated between the buffalo and cattle populations (respectively 1.9 $\pm$ 0.97 and 1.18 $\pm$ 0.69) and the transmission rate of 0.35 $\pm$ 0.35 from the impala to the buffalo population.

Using the "mugration" model, we determined numerous transmission events between buffalo and cattle populations and lower numbers of transmission events between the impala and cattle populations (see Figure 3a). In comparison, using BASTA, we observed few transmission events from the cattle to the buffalo populations but more transmission events between the impala and cattle populations (see Figure 3b).

a. FMDV SAT1: "mugration" model

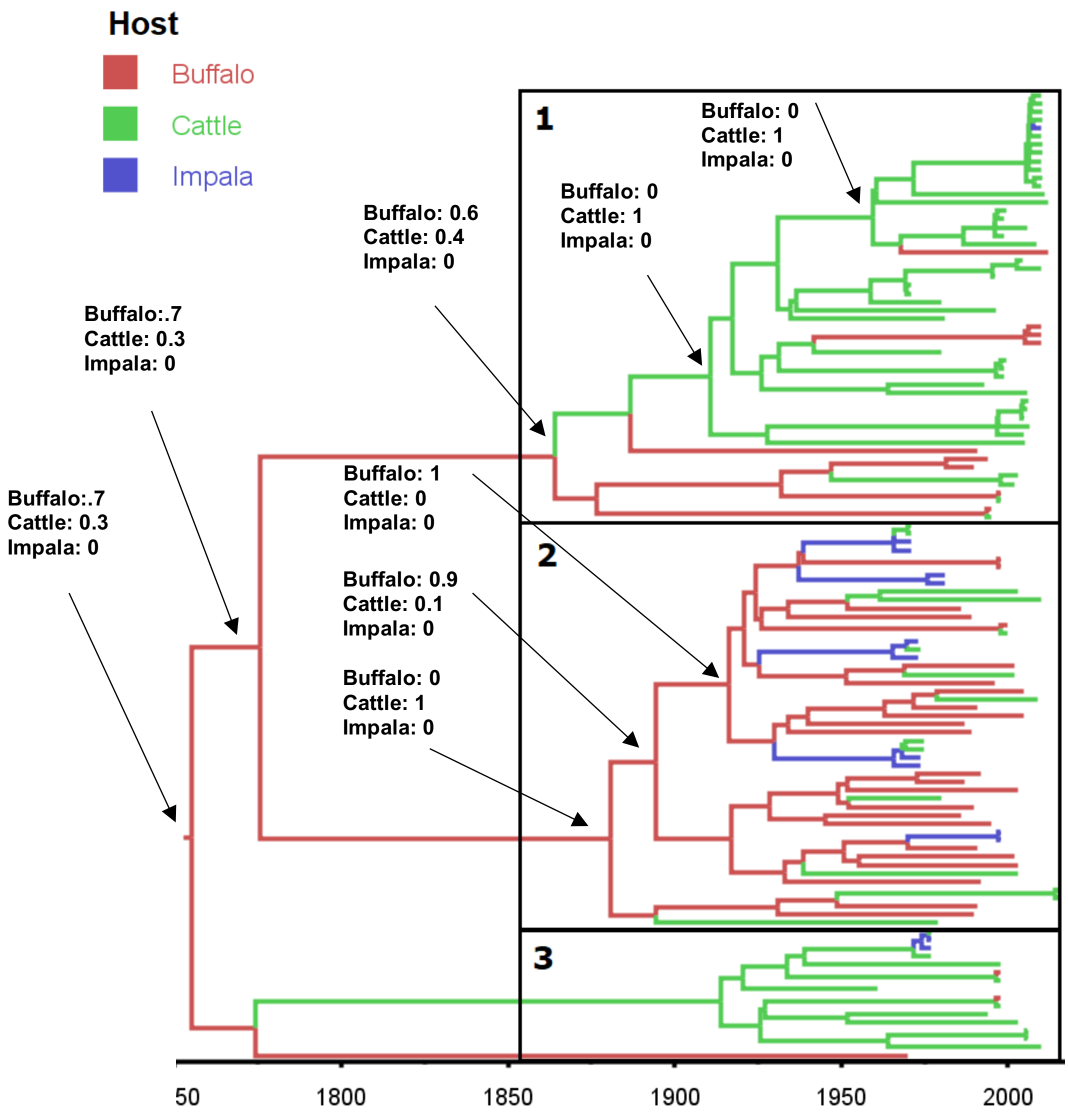

b. FMDV SAT1: BASTA model

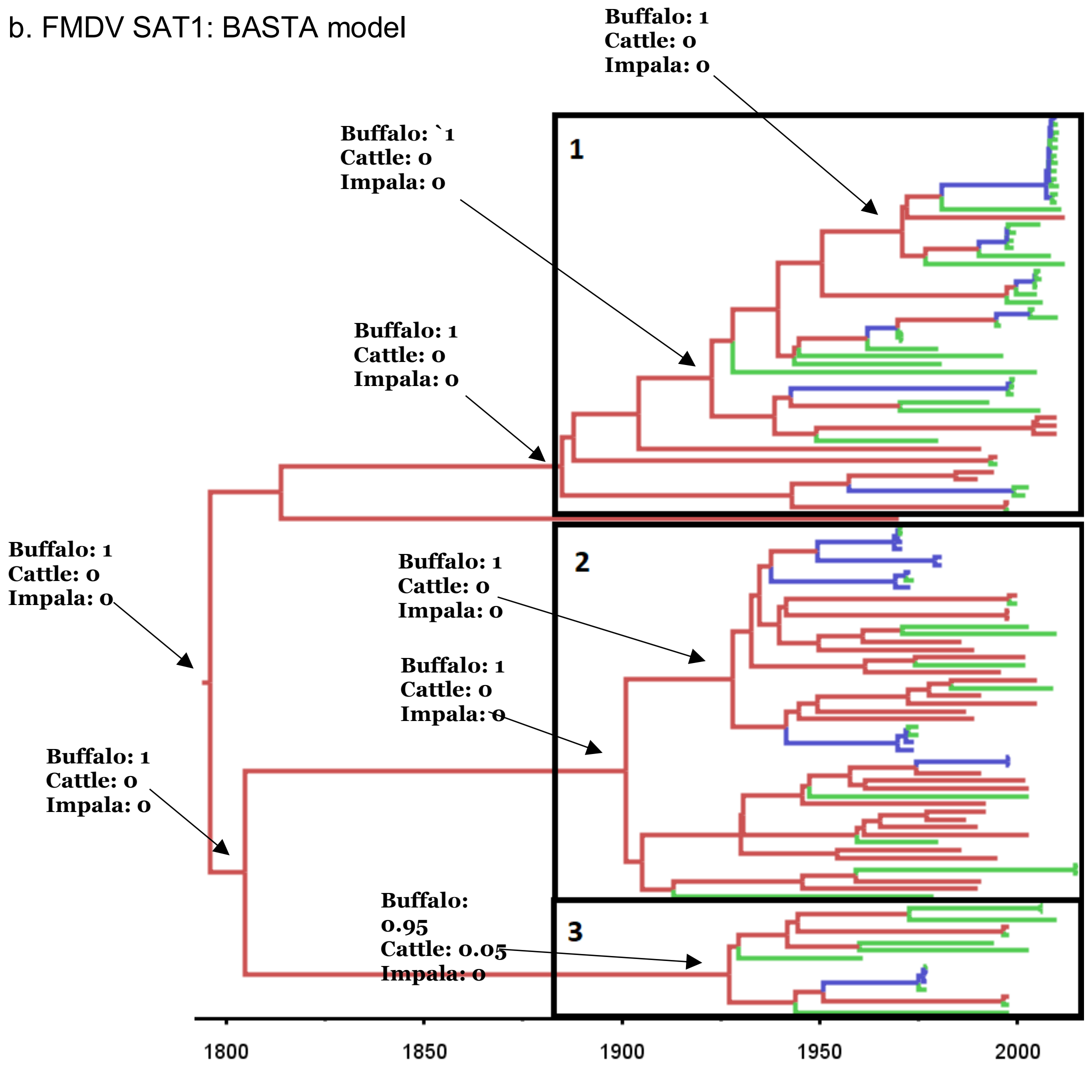

c. FMDV SAT1: MASCOT model

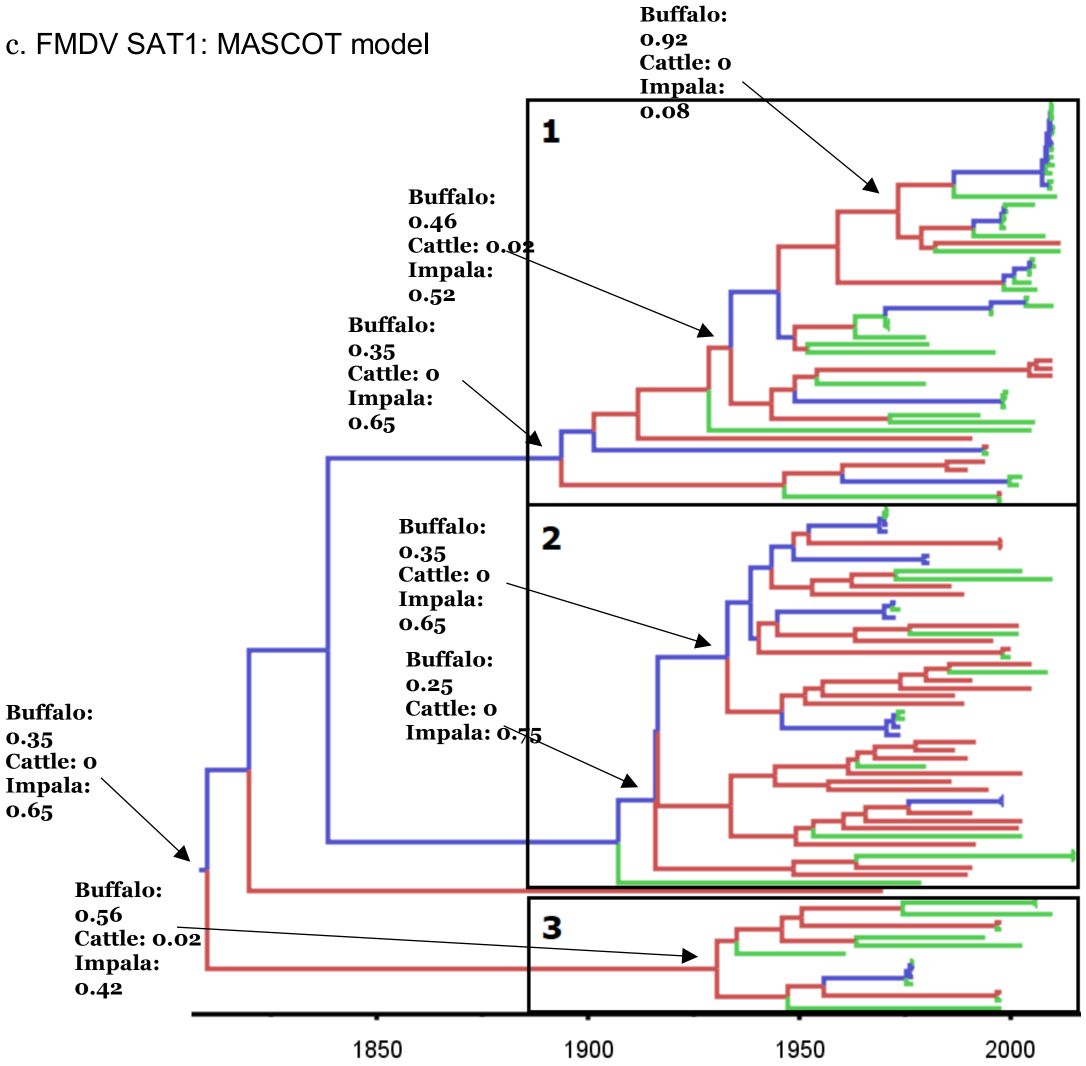


**Figure 1. Bayesian MCC time scaled discrete phylogeographic tree for the serotype SAT1 using 113 VP1 sequences.**

a. Phylogenetic tree estimated using the "mugration" approach implemented in BEAST b. Phylogenetic tree estimated using the BASTA approach implemented in BEAST2. c. Phylogenetic tree estimated using the MASCOT approach implemented in BEAST2. The phylogeny branches are coloured according to their descendent nodes host with the key for colours shown on the upper left of the figure. The identified clades were isolated and numerated. Specific nodes of the trees were annotated with hosts posterior probabilities.

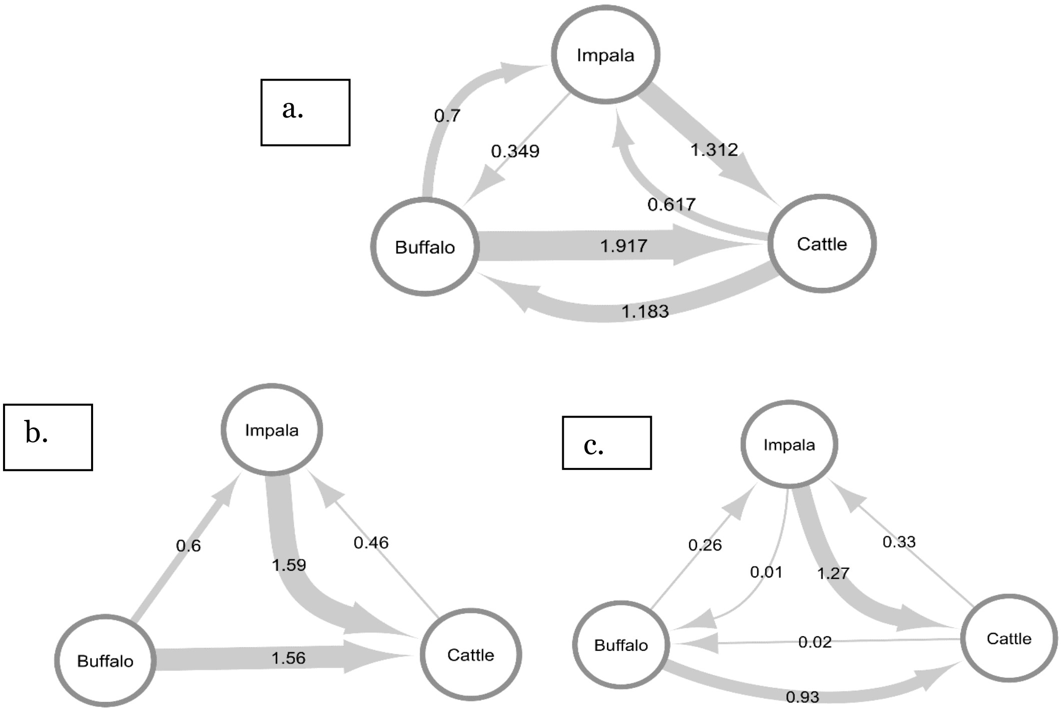


**Figure 2. Estimated transmission rates between the three potential hosts for the SAT1 FMDV serotype**. a. Using the “Mugration” approach in BEAST. b. Using the BASTAapproach in BEAST2. c. Using the MASCOT approach in BEAST2.

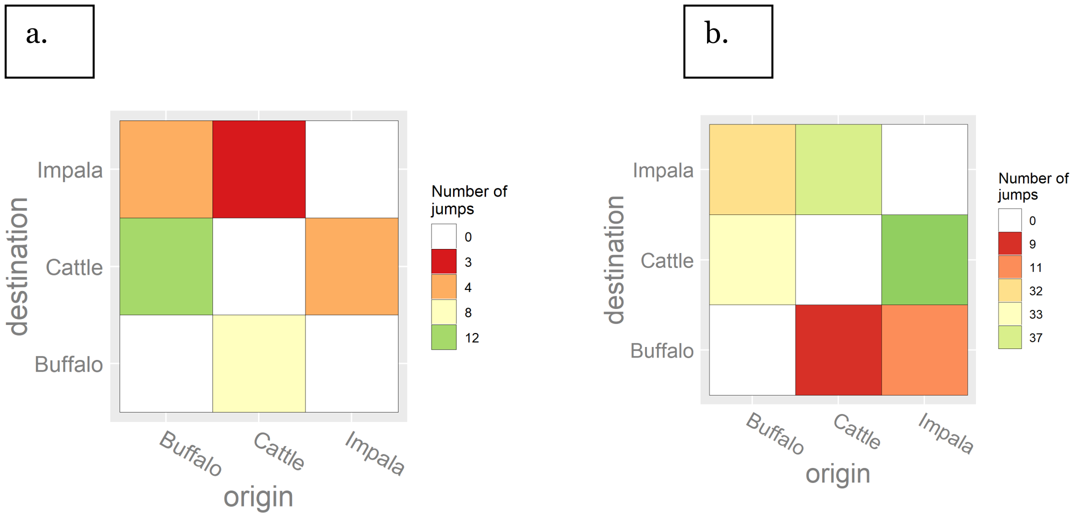


**Figure 3. Heatmap showing the number of transitions between the sampled hosts for theSAT1 FMDV serotype obtained through a Markov jump analysis.** The heatmaps are coloured according to the number of estimated transitions between hosts. a. Host transition using the ”mugration” model b. Host transition using the BASTA model.

When analyzing the FMDV SAT2 serotype dataset we estimated a similar clock rate of $1.1x10^{-3}$ substitution/site/year between the tree models. However, we estimated a longer tree height of 430 years for the "mugration" method than for the two structural coalescent approximation methods that had a tree height closer to 350 years. Both BASTA and MASCOT models estimated a similar buffalo population size (around 1200 for both models) and impala population size (2/3 in both models) volved in the SAT2 serotype spread. However, the BASTA model estimated a slightly larger cattle population (3.9 $\pm$ 1.89) than MASCOT (1.8 $\pm$ 1) (see Supplementary table **S1** and Supplementary table **S2)**

With all three methods we isolated the same fives main clades. Because the "mugration" model estimated that two out of five clades composing the phylogenetic tree of SAT2 were entirely composed of cattle nodes (clades two and five), it estimated an important role of the cattle population in the transmission of the virus (see Figure 4a). However, for this serotype we cannot observe transmission events from the impala toward the cattle populations. In comparison, with most of the tree composed of buffalo nodes, the BASTA and MASCOT models estimate a less important role for the cattle population in the transmission of FMDV SAT2 serotype (see Figure 4b and Figure 4c.).

Additionally, in these two models we estimated multiple transmission events of FMDV SAT2 serotypes from buffalo to impala and then to the cattle population. Again, as with the FMDV SAT1 serotype, the estimated number of transitions and transmission rates between the different populations estimated for the SAT2 serotype stresses the differences between the "mugration" model and both structural coalescent approximation approaches.

In both the BASTA and MASCOT models, we estimated low transmission rates (less than 1-2) from the impala and cattle populations toward the buffalo population (see Figure 5b and Figure 5c). In both approaches, we estimated higher rates of transmission from buffaloes to impalas (1 $\pm$ 0.44 in BASTA and in 0.466 $\pm$ 0.28 in MASCOT) than from cattle to impalas (0.24 $\pm$ 0.31 in BASTA and 0.25 $\pm$ 0.33 in MASCOT). By looking at the transmission rate, the main difference between the two structural coalescent models is that the BASTA model estimated a higher transmission rate from impalas to cattle than from buffaloes to cattle (1.38 $\pm$ 0.77 and 0.7 $\pm$ 0.46), while the MASCOT model estimated similar transmission rates (0.95 $\pm$ 0.66 and 0.96 $\pm$ 0.43). In comparison to the structural coalescent approximation approaches, most of the transmission rates obtained with the "mugration" model were between 0.5 and 1, except for the transmission rate from the buffalo to the cattle population which is equal to 2.8 $\pm$ 1.44 (see Figure 5a).

The "mugration" model estimated that multiple transmission events occurred between the buffalo and cattle populations and that no transmission events occurred between the impala and cattle populations (see Figure 6a). In comparison, the BASTA approach estimated multiple transmission events from the impala and buffalo populations toward the cattle population and a small number of transmissions with the cattle population as origin (see Figure 6b).

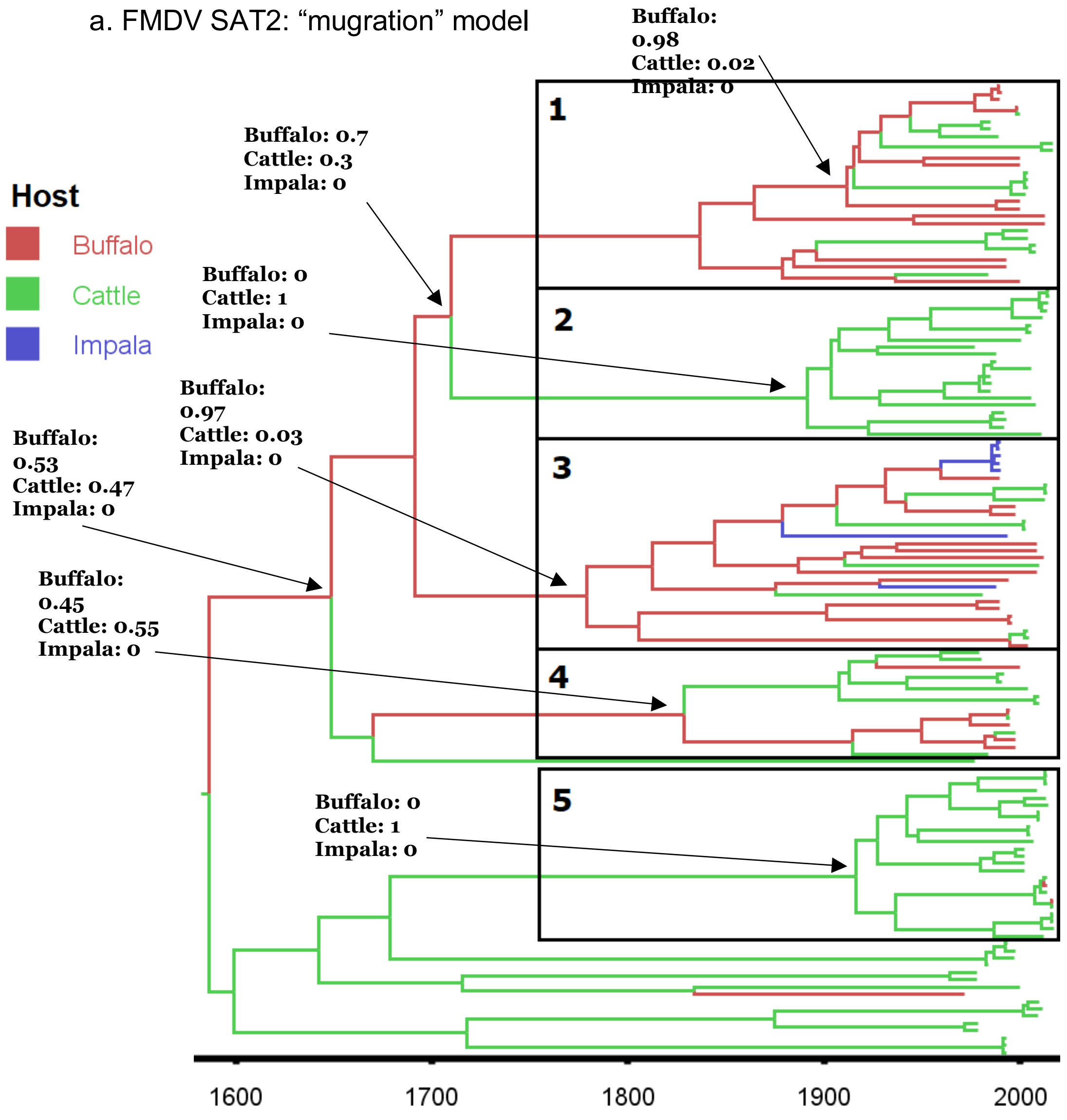

b. FMDV SAT2: BASTA model

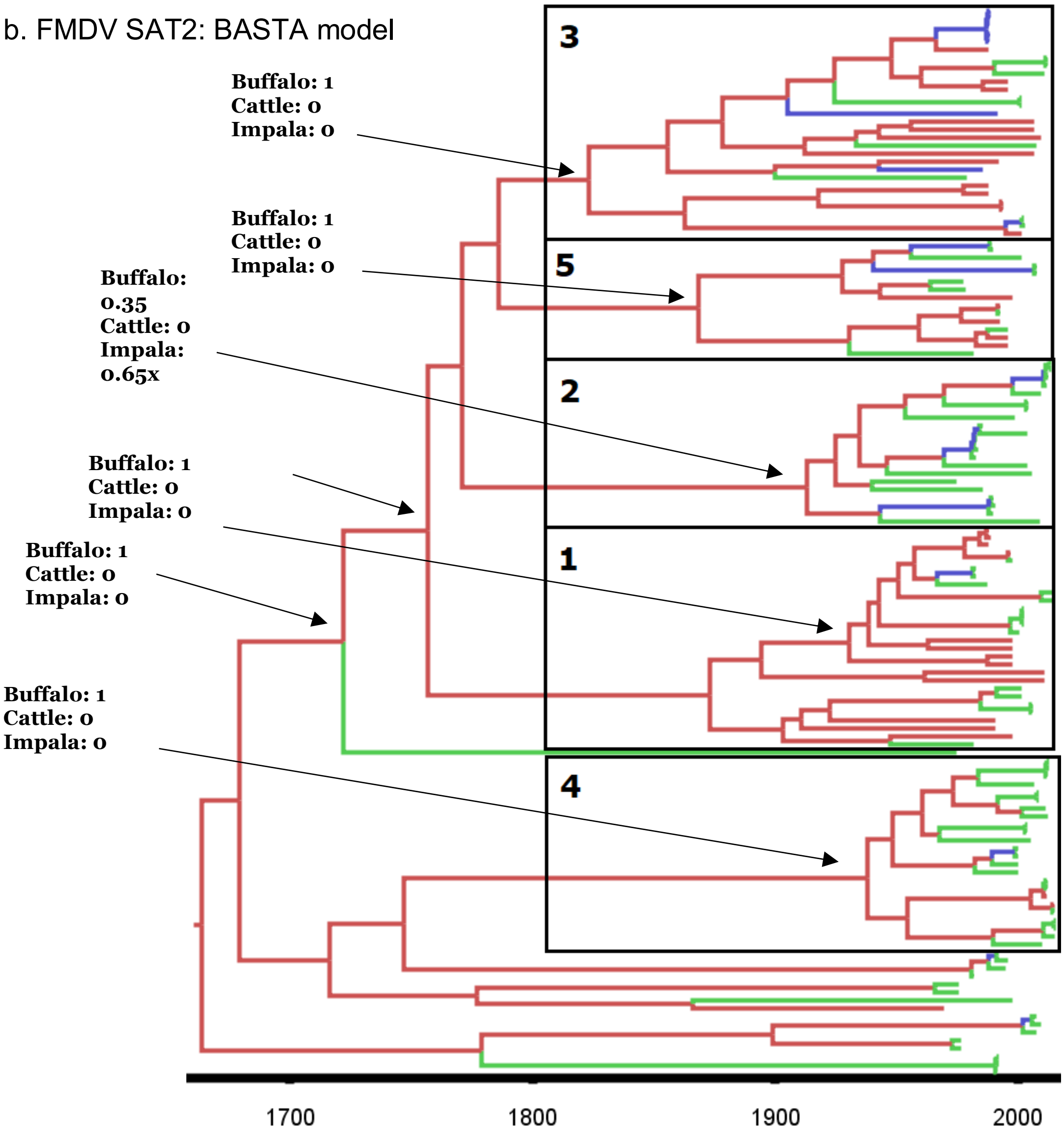

c. FMDV SAT2: MASCOT model

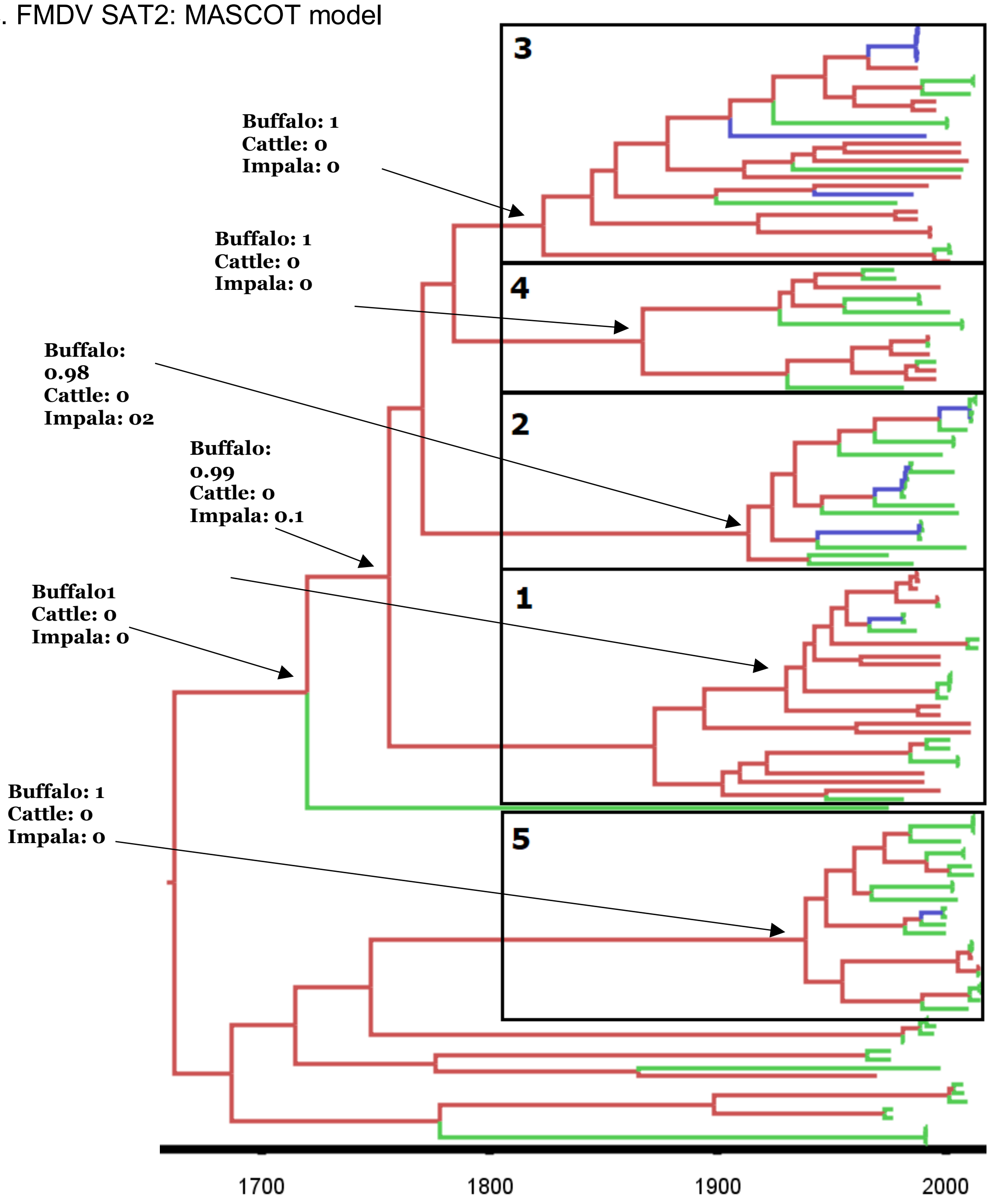


**Figure 4. Bayesian MCC time scaled discrete phylogeographic tree for the serotype SAT2using 135 VP1 sequences**. a. Phylogenetic tree estimated using the “mugration” approach implemented in BEAST b. Phylogenetic tree estimated using the BASTA approach implemented in BEAST2. c. Phylogenetic tree estimated using the MASCOTapproach implemented in BEAST2. The phylogeny branches are coloured according totheir descendent nodes host with the key for colours shown on the upper left of the figure. The identified clades were isolated and numerated. Specific nodes of the trees were annotated with hosts posterior probabilities

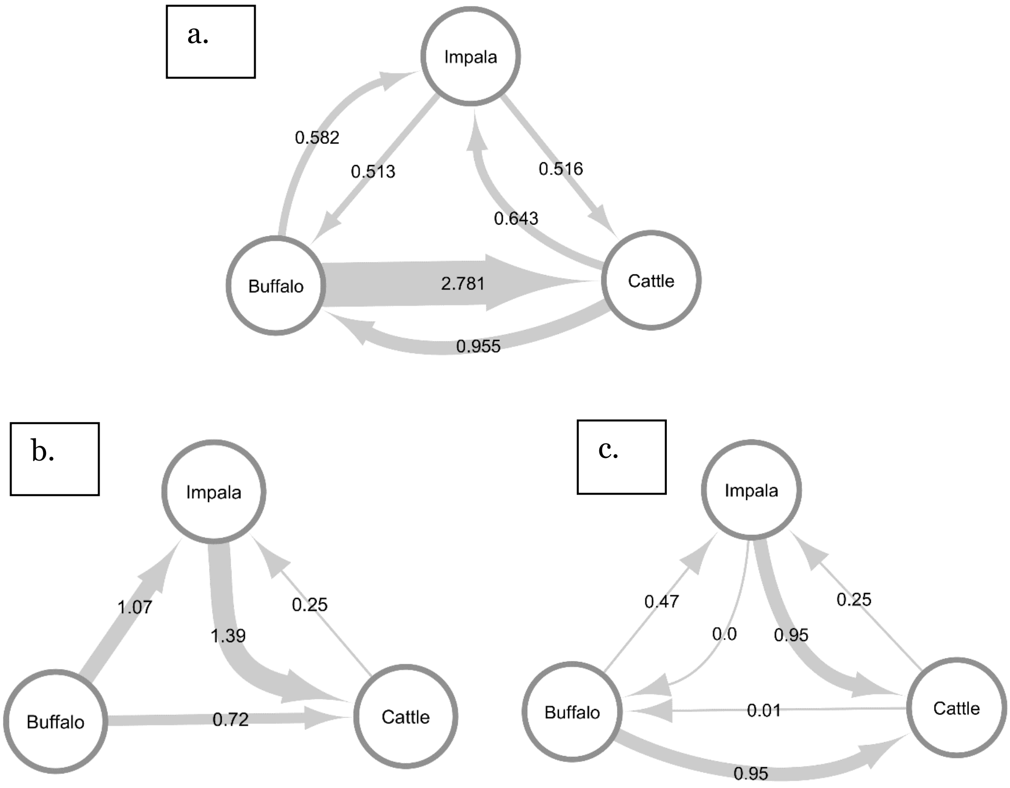


**Figure 5. Estimated transmission rates between the three potential hosts for the SAT2 FMDV serotype.** a. Using the "Mugration" approach in BEAST. b. Using the BASTAapproach in BEAST2. c. Using the MASCOT approach in BAST2.

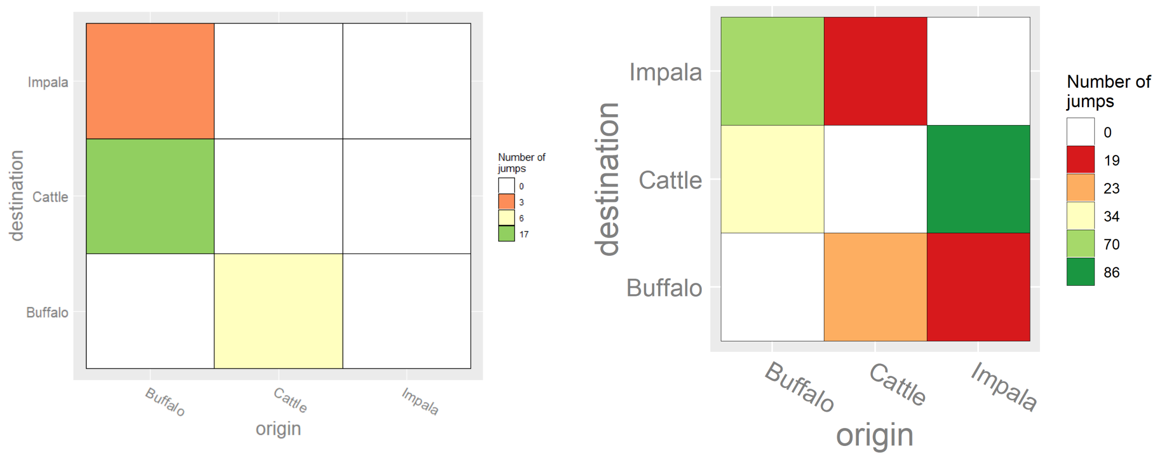


**Figure 6. Heatmap showing the number of transitions between the sampled hosts for theSAT2 FMDV serotype.** The heatmaps are colored according to the number of estimated transitions between hosts. a. Host transition using the "mugration" model b.Host transition using the BASTA model.
